# *De novo* origins of multicellularity in response to predation

**DOI:** 10.1101/247361

**Authors:** Matthew D. Herron, Joshua M. Borin, Jacob C. Boswell, Jillian Walker, I-Chen Kimberly Chen, Charles A. Knox, Margrethe Boyd, Frank Rosenzweig, William C. Ratcliff

## Abstract

The transition from unicellular to multicellular life was one of a few major events in the history of life that created new opportunities for more complex biological systems to evolve. Predation is hypothesized as one selective pressure that may have driven the evolution of multicellularity. Here we show that *de novo* origins of simple multicellularity can evolve in response to predation. We subjected outcrossed populations of the unicellular green alga *Chlamydomonas reinhardtii* to selection by the filter-feeding predator *Paramecium tetraurelia*. Two of five experimental populations evolved multicellular structures not observed in unselected control populations within ~750 asexual generations. Considerable variation exists in the evolved multicellular life cycles, with both cell number and propagule size varying among isolates. Survival assays show that evolved multicellular traits provide effective protection against predation. These results support the hypothesis that selection imposed by predators may have played a role in some origins of multicellularity.

---

Nearly all macroscopic life is multicellular. As Leo Buss emphasized in *The Evolution of Individuality*, the very existence of integrated multicellular organisms is an outcome of evolutionary processes, not a starting condition^1^. It seems, in fact, to be a common outcome: multicellular organisms have evolved from unicellular ancestors dozens of times^2–4^. Animals, land plants, fungi, red algae, brown algae, several groups of green algae, cellular and acrasid slime molds, and colonial ciliates, among others, each descend from a different unicellular ancestor^4,5^.

Two retrospective approaches, comparative methods and the fossil record, have proven valuable in reconstructing how these transitions may have occurred. Although both approaches have been critical to our understanding of early multicellular evolution, each has its limitations. For most multicellular groups, little or no fossil evidence exists that is relevant to the first steps in the transition from unicellular to multicellular life. Comparative methods suffer from a lack of intermediate forms between the multicellular organisms we are interested in and their extant unicellular relatives. Furthermore, direct knowledge of unicellular ancestors is not available. Extant unicellular relatives often serve as stand-ins, but this is a poor approximation, as they have been evolving independently as single-celled organisms since they diverged from their multicellular relatives.

A third, prospective, approach designed to circumvent these limitations has emerged in the last decade. The experimental evolution of multicellularity in otherwise unicellular microbes enables real-time observations of morphological, developmental, and genetic changes that attend the transition to multicellular life. Boraas and colleagues exposed cultures of the green alga *Chlorella vulgaris* to predation by the flagellate *Ochromonas vallescia*, resulting in the evolution of small, heritably stable algal colonies^6^. Becks and colleagues showed that exposure to the predatory rotifer *Brachionus calyciflorus* selected for heritable changes in the rate of formation of multicellular palmelloids in the green alga *Chlamydomonas reinhardtii*^7^. Ratcliff and colleagues have shown that selection for an increased rate of settling out of liquid suspension consistently results in the evolution of multicellular ‘snowflake’ colonies in the yeast *Saccharomyces cerevisiae*^8^ and also results in the evolution of simple multicellular structures in *C. reinhardtii*^9^.

Predation has long been hypothesized as a cause for the evolution of multicellularity, as most predators can only consume prey within a narrow range of sizes^2,10–12^. Filter-feeding predators are common in aquatic ecosystems, and algae that are larger than a threshold size are largely immune to them^10^. Thus, predation is an ecologically plausible selective pressure that could explain at least some origins of multicellularity.

In this study, we present experiments in which we used the ciliate predator *Paramecium tetraurelia* to select for the *de novo* evolution of multicellularity in outcrossed populations of *C. reinhardtii*. We describe the heritable multicellular life cycles that evolved and compare them to the ancestral, unicellular life cycle. In addition, we show that the evolved multicellular life cycles are stable over thousands of asexual generations in the absence of predators. Comparative assays show that the evolved multicellular phenotypes provide a fitness advantage over its unicellular ancestor in the presence of predators. Because *C. reinhardtii* has no multicellular ancestors, these experiments represent a completely novel origin of obligate multicellularity^13,14^.

## Results

After 50 weekly transfers (~750 generations), simple multicellular structures evolved in two of five predator-selected populations (B2 and B5). Such multicellular structures were not observed in any of the control populations. Eight strains were isolated from each of three populations (B2, B5, K1). We focused our analyses on five focal strains from B2 (B2-01, B2-03, B2-04, B2-10, B2-11) and two strains from B5 (B5-05, B5-06). Of the isolates from the control population that evolved in the absence of predators (K1), we analyzed two strains (K1-01, K1-06). Phenotypes of other isolates from populations B2, B5 and K1 did not differ qualitatively from the focal strains and were not investigated further. The strains have maintained their evolved characteristics of simple multicellularity in the absence of predators for four years as unfrozen, in-use laboratory strains. Therefore, we are confident that the phenotypic traits that we report below are stably heritable.

Some strains, notably those from population B5, commonly formed stereotypic eight-celled clusters, with an apparent unicellular and tetrad life stage (Fig. 1B). Other strains, notably those from population B2, appeared to form amorphous clusters of variable cell number (Fig. 1A). Other phenotypic differences could be easily discerned by light microscopy. For example, in Fig. 1, an external membrane is visible around both evolved multicellular colonies, indicating that they formed clonally via repeated cell division within the cluster, rather than via aggregation.

**Figure 1.**
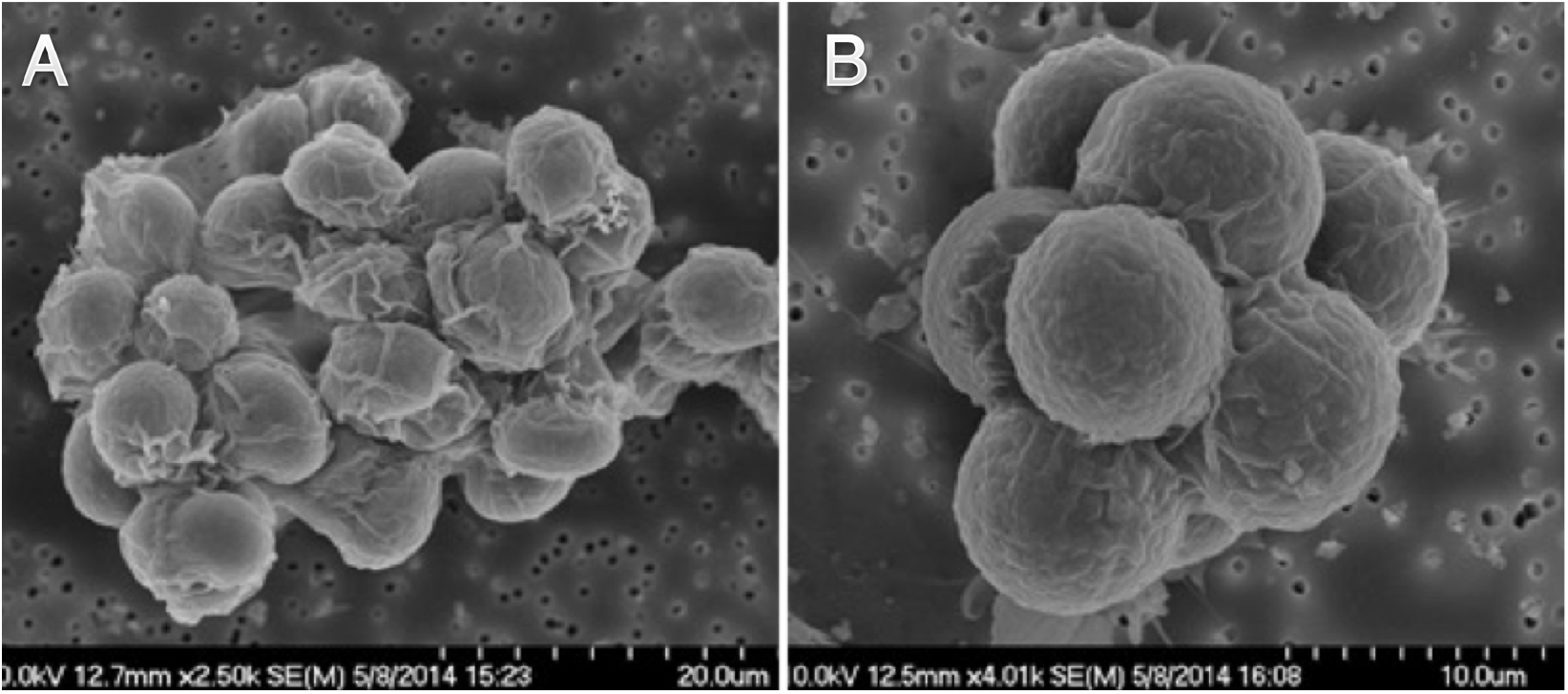
Scanning electron micrographs of representative multicellular colonies from evolved populations. [A] shows an amorphous cluster from population B2. Cell number varies greatly between clusters in this clone and between clones in this population. [B] shows an eight-celled cluster from population B5. Octads were frequently observed in both populations.

We found a variety of life cycles within and among populations. Using time-lapse videos of evolved strains growing in liquid culture in 96-well tissue culture plates, we qualitatively classified strains into four life cycle categories to compare their similarities and differences (Fig. 2A-D). We discuss each life cycle category below, and then present quantitative measurements on parent and propagule cluster sizes.

**Figure 2.**
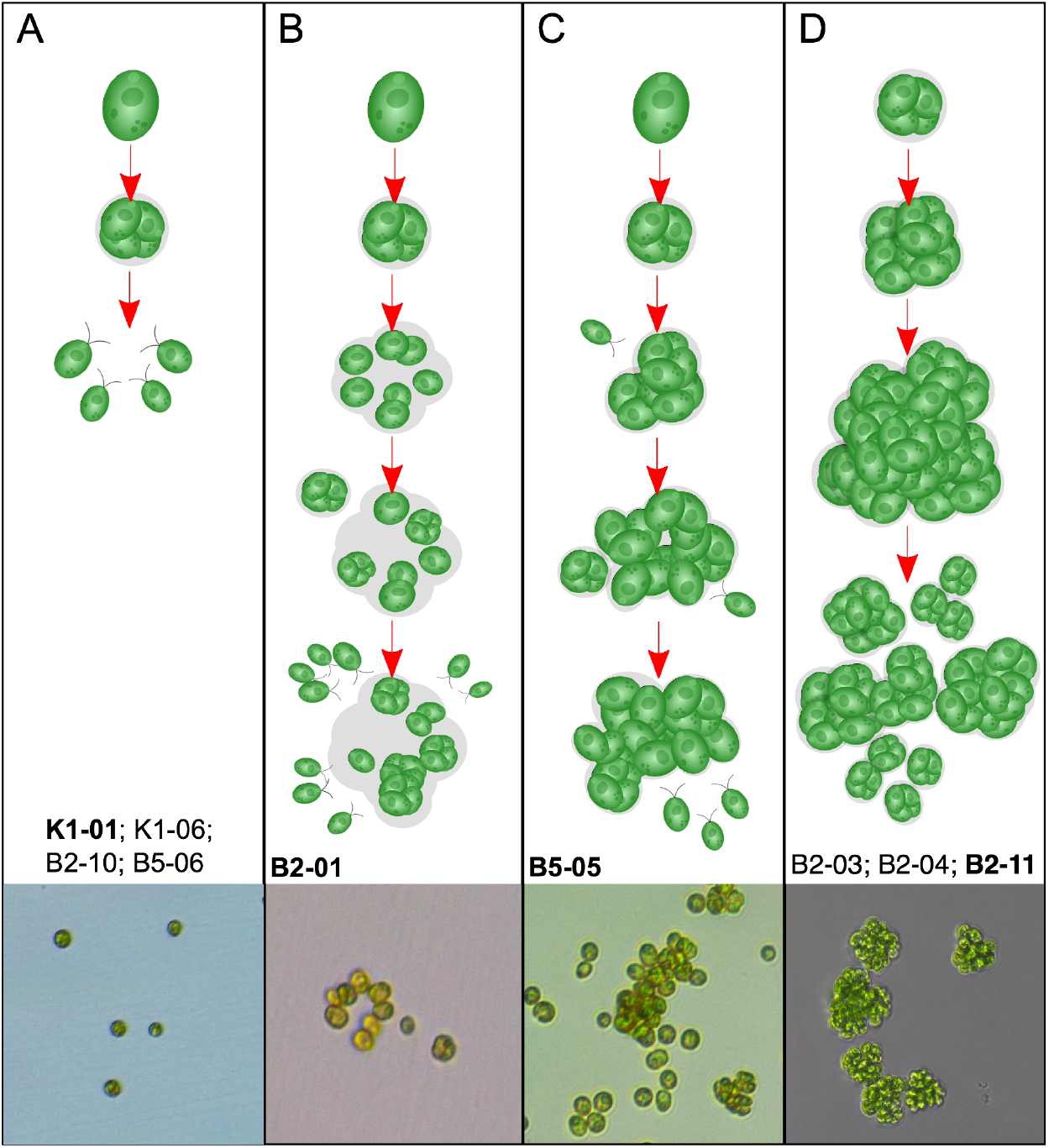
Depiction of *C. reinhardtii* life cycles following evolution with (B2, B5) or without (Kl) predators for 50 weeks. Categories (A-D) show a variety of life cycle characteristics, from unicellular to various multicellular forms. Evolved strains were qualitatively categorized based on growth during 72-hour time-lapse videos. Strains within each life cycle category are listed below illustrations. Representative microscopic images of each life cycle category are at the bottom (Depicted strain in **boldface**).

In population K1, which evolved without predators, ancestral life cycle characteristics of the unicellular, wild-type *C. reinhardtii* were retained (Fig. 2A, Supplemental Videos 1 and 2). Specifically, as cells reproduce asexually, they lose motility and undergo 2-5 rounds of mitosis before releasing motile, single-celled propagules. It should be noted that even in wild-type *C reinhardtii*, the dividing parent cluster is a *transient* multicellular stage; however, it does not persist after propagules are released. Interestingly, in the two populations that evolved multicellularity in response to predation (B2, B5), strains B5-06 and B2-10 retained a life cycle typical of the ancestral, wild-type *C. reinhardtii* (Supplemental Videos 3 and 4, respectively).

Life cycles of the remaining strains isolated from populations B2 and B5 were distinct from the ancestor, as clusters of various sizes persist through multiple rounds of reproduction. Ordinarily, strain B2-01 releases motile, single-celled propagules during reproduction, similar to the ancestor (Supplemental Video 5). However, in some clusters, cells undergoing division separate, but remain proximately located because they are embedded in an extracellular matrix (ECM) of the parent cluster (Fig. 2B). As these cells continue to grow and divide, some remain embedded in the ECM, which creates growing aggregations of cells. Strain B5-05 also produces motile, single-celled propagules that are often embedded in the maternal ECM (Supplemental Video 6). In addition to retaining propagules embedded in the ECM, growing clusters of B5-05 ensnare free-swimming cells, creating aggregations that grow much larger than those of B2-01 (Fig. 2C).

Conversely, three of the strains isolated from population B2 exist in cell clusters comprised only of direct descendants, as opposed to chimeric aggregations with free-swimming cells. Clusters from strains B2-11, B2-03, and B2-04 grow in tightly associated groups of direct descendants embedded in the maternal cell wall (Fig. 2D; Supplemental Videos 7, 8, and 9, respectively). Development in these isolates is therefore strictly clonal, with important implications for evolvability. Since the cells within a multicellular structure are likely to be genetically identical, other than differences resulting from new mutations, genetic variation in a population would be partitioned primarily among colonies. The clonal development observed in these isolates therefore suggests that the observed multicellular clusters would be well-suited to serving as units of selection.

In key respects, the isolates from population B2 appear to have recapitulated early steps in volvocine evolution; in fact, the evolved multicellular algae are similar in their gross morphology to small colonial volvocine algae such as *Pandorina*. Furthermore, a degree of genetic control of cell number similar to that seen in undifferentiated colonial volvocine algae manifests in our evolved multicellular strains. Propagules of these strains are typically multicellular and, critically, no motile propagules were observed in these strains. Propagules from the evolved multicellular strains were nearly all 4-, 8-, or 16-celled, a range similar to that of small colonial volvocine algae and smaller than that of *Pleodorina starrii*, in which propagules of a single genotype can span a 16-fold range of cell numbers^15^.

In order to determine the size attained by evolved strains, we sampled, stained, and imaged growing cultures in tubes every 12 hours over a 6-day period. We find that cluster sizes of strains that evolved without predators (K1-01, K1-06) and strains that retained ancestral life cycle traits, despite evolving with predators (B5-06, B2-10), remain small throughout the six days of growth (Fig. 3). Even at their largest, median cluster sizes of replicate populations of strains K1-01 and K1-06 were all one cell per cluster, showing that these strains exist as unicellular individuals with a transient a multicellular stage occurring during reproduction. Similarly, we find that cells of strain B2-01 exist primarily as single cells, with median sizes averaging 1.8 cells. Under time-lapse observation, we found that dividing cells of B2-01 would occasionally separate from direct contact with each other while still being held together by maternal ECM/cell wall. This was seen as white spaces appearing between cells, which then remained equidistant in a rigid structure. During the procedures to stain and image cells, it is possible that these semi-separated cells were dispersed, leading to a slightly lower estimate of median cluster sizes. The median cluster sizes of replicate populations of strain B5-05 averaged 4.0 cells, larger than strains that retained the ancestral phenotype as well as B2-01. Unlike strains B2-01 and B5-05, in which ECM holds cells in clusters, cells of the largest strains are visibly encapsulated within the mother cell wall. Medians of replicate populations of these strains (B2-11, B2-03, and B2-04) averaged 6.2, 5.6, and 5.6, respectively.

**Figure 3.**
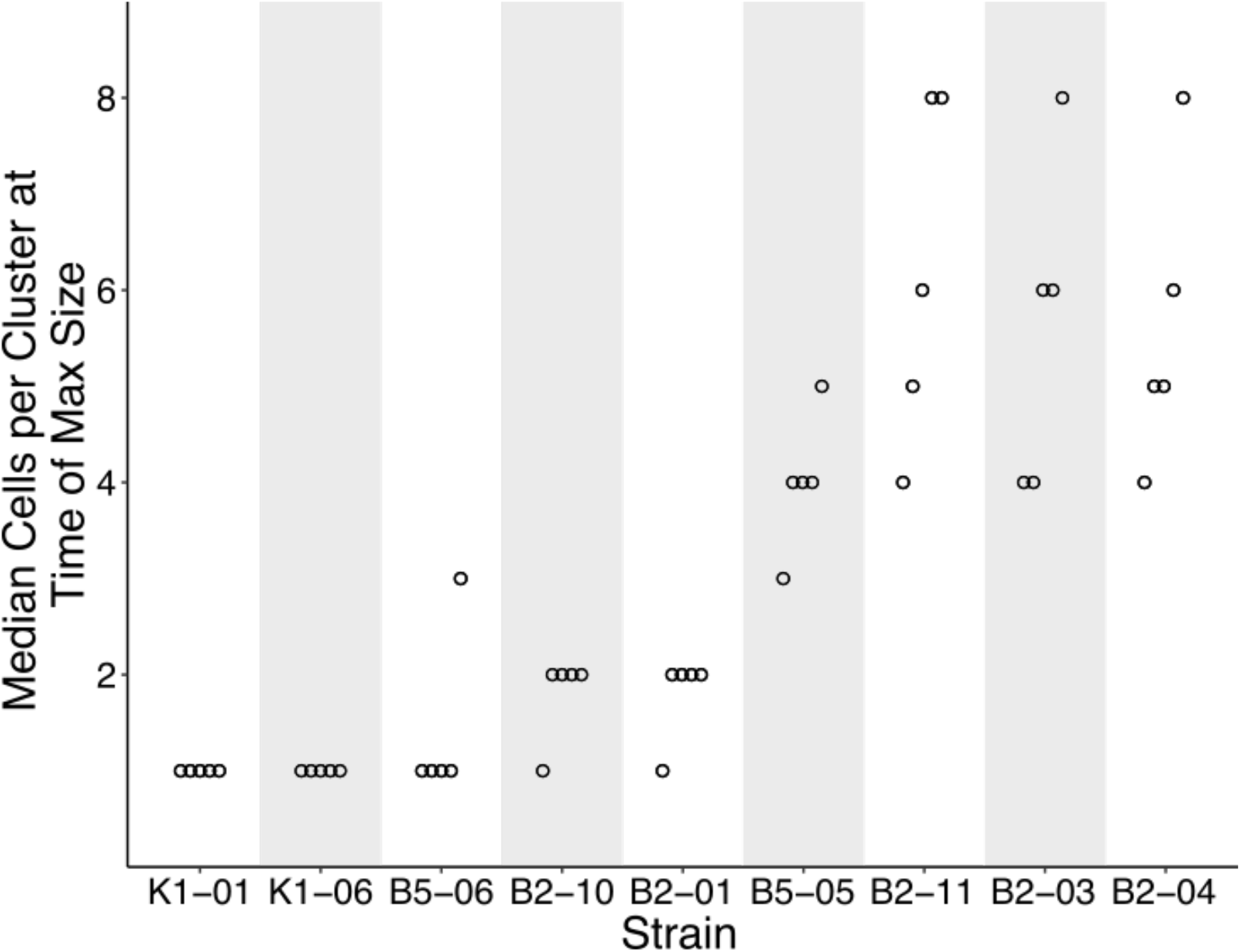
Median cluster sizes of evolved strains at the time-point where strains reached maximum size. To determine the time where strains were largest, we calculated the mean from five replicate medians (open dots), for each strain and time-point. Because data are not normally distributed, medians were chosen to approximate central tendency. Cells per cluster were measured by sampling strains over six days of growth, staining nuclei with DAPI, and imaging using fluorescent microscopy. From left to right, time-points of maximum size for each strain were: 12, 12, 72, 120, 96, 108, 84, 72, 72, 96, 96, and 72 hours. From left to right, median sizes of replicate populations averaged 1, 1, 1.4, 1.8, 1.8, 4, 6.2, 5.6, and 5.6 cells per cluster. Sizes at the initial time-point (0 hrs.) were omitted from analysis because they represent starting conditions. Shading is only for ease of visualization.

Because cells were mixed several times during the staining and imaging process, it is possible that some clusters of cells were disrupted. Thus, cluster sizes from fluorescent microscopy analysis are likely underestimates of cluster size, which may or may not affect cluster size estimates for certain strains (i.e., those held together by extracellular matrix rather than the maternal cell wall) more than others.

In order to determine propagule sizes of evolved strains, we manually analyzed 12-hour time-lapse videos and recorded the numbers of cells in propagules released from parent clusters. For most strains the majority of propagules were single-celled, as observations were skewed toward smaller propagules, which require less biomass to produce (Fig. 4). To accurately depict the context of individual cells in propagules, we also show the data as boxplots, weighted by biomass (Fig. 4, black boxplots). Strains that maintained ancestral characteristics and had the smallest average maximum cluster sizes (K1-01, K1-06, B5-06, B2-10) released almost exclusively single-celled propagules. Although cells from strains B2-01 and B5-05 often exist in clusters, the vast majority of propagules released by these reproducing clusters were single-celled as well. In strains with larger average maximum cluster sizes (B2-11, B2-03, B2-04), cells are more frequently released in multicellular propagules of up to 64 cells in size. Thus, among the strains that evolved under predation, a variety of size-related traits emerged in both cluster size and propagule size.

**Figure 4.**
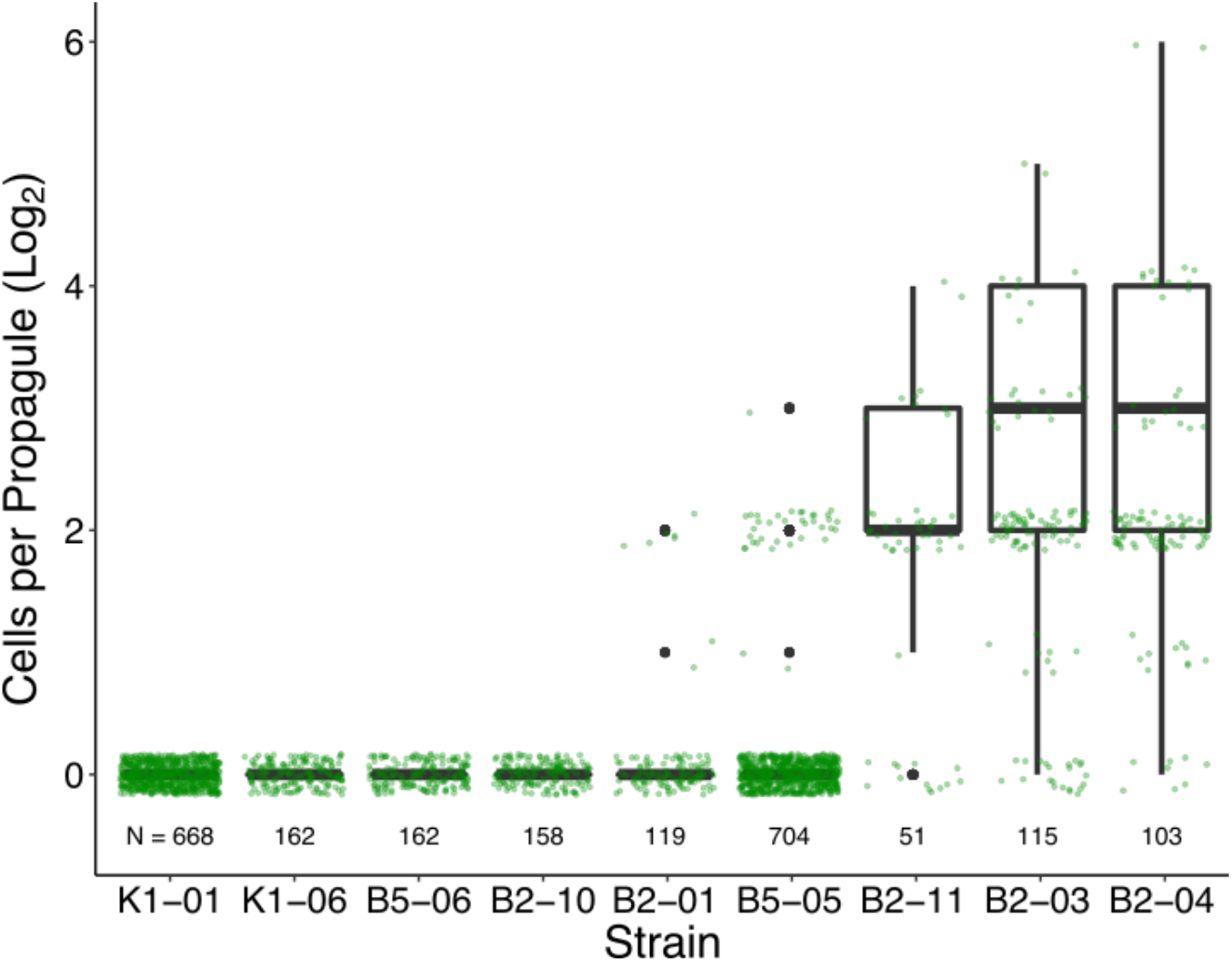
Sizes of propagules released by evolved strains during a 12-hour time-lapse. Propagule sizes were manually measured from time-lapse videos. Green dots show actual observations of propagule size. Because the frequency of observations is skewed toward smaller propagule sizes, we show the data as weighted boxplots as well, where a 32-celled propagule has 32 times the weight of a single-celled propagule. Sample sizes of propagule observations for each strain are indicated along the bottom of the plot. Strains are ordered by weighted-mean propagule size.

Multicellularity appears to provide an effective defense against predation. Comparisons of predation rates were made by measuring the absorbance of each of the four replicates without predators and dividing by the average of the four replicates with predators. This value, labeled Δabs, shows markedly different trajectories for unicellular and multicellular treatments with predators (Fig. 5). The unicellular strains, on average, experienced ~2.5-fold greater rates of predation compared to the multicellular strains. That is, the mean Δabs at hour 117 (day 5) were 0.367 and 0.140 for unicellular and multicellular strains, respectively. To simplify the analysis, inferences were drawn from the final time-point measurements (hour 117). A test for an effect of phenotype (unicellular or multicellular) on Δ_abs_ values shows that phenotype has a strong effect on algal density (ANOVA, F_1.22_ = 8.65, P = 0.0076).

**Figure 5.**
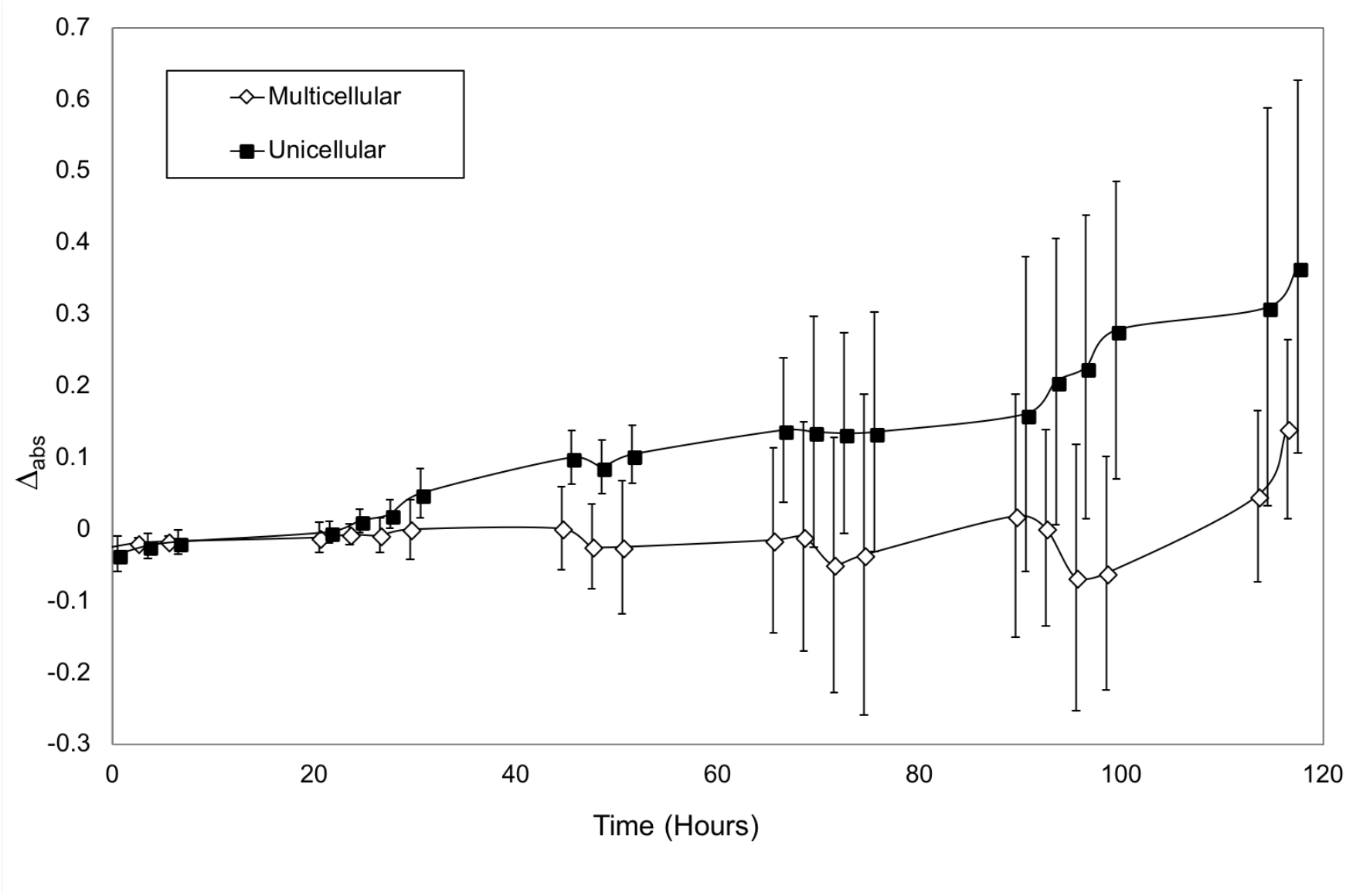
Average absorbance value differences (Δ_abs_) between populations with and without predation for evolved multicellular and unicellular *C. reinhardtii* strains under predation. Multicellular strains averaged a much lower change in absorbance over the duration of the experiment than unicellular strains. Error bars are standard deviations of twelve strains per treatment. Markers are offset 0.5 h so that error bars can be seen.

## Discussion

Our results show that the transition to a simple multicellular life cycle can happen rapidly in response to an ecologically relevant selective pressure. By increasing in size beyond the “predation threshold”^16^ of a filter-feeding predator, multicellular *C. reinhardtii* (evolved from an ancestrally unicellular lineage) are protected from predation for at least part of their life cycle. Under selection for increased size, formation of multicellular structures may be an easier route than increasing cell size because of trade-offs imposed by scaling relationships (primarily the reduction in surface-area-to-volume ratio), because more mutational paths are available, and/or because available mutations have fewer or less severe pleiotropic effects.

These results support the view that predation may have played a role in at least some origins of multicellularity. Consistent with previous results^6–9^, predation can drive the evolution of simple multicellular structures, and these structures provide protection from predators^6,17^. The ‘chicken-and-egg’ problem brought up by some authors – that multicellular predators are required to drive the evolution of multicellularitye.g.,^2,12^ – does not exist: unicellular filter-feeding protists exist, including, for example, *Paramecium*, and there is no reason to think that such predators only arose after animals. Furthermore, some origins of multicellularity, including those of brown and volvocine algae, postdate that of animals, and could therefore have been driven by animal predation.

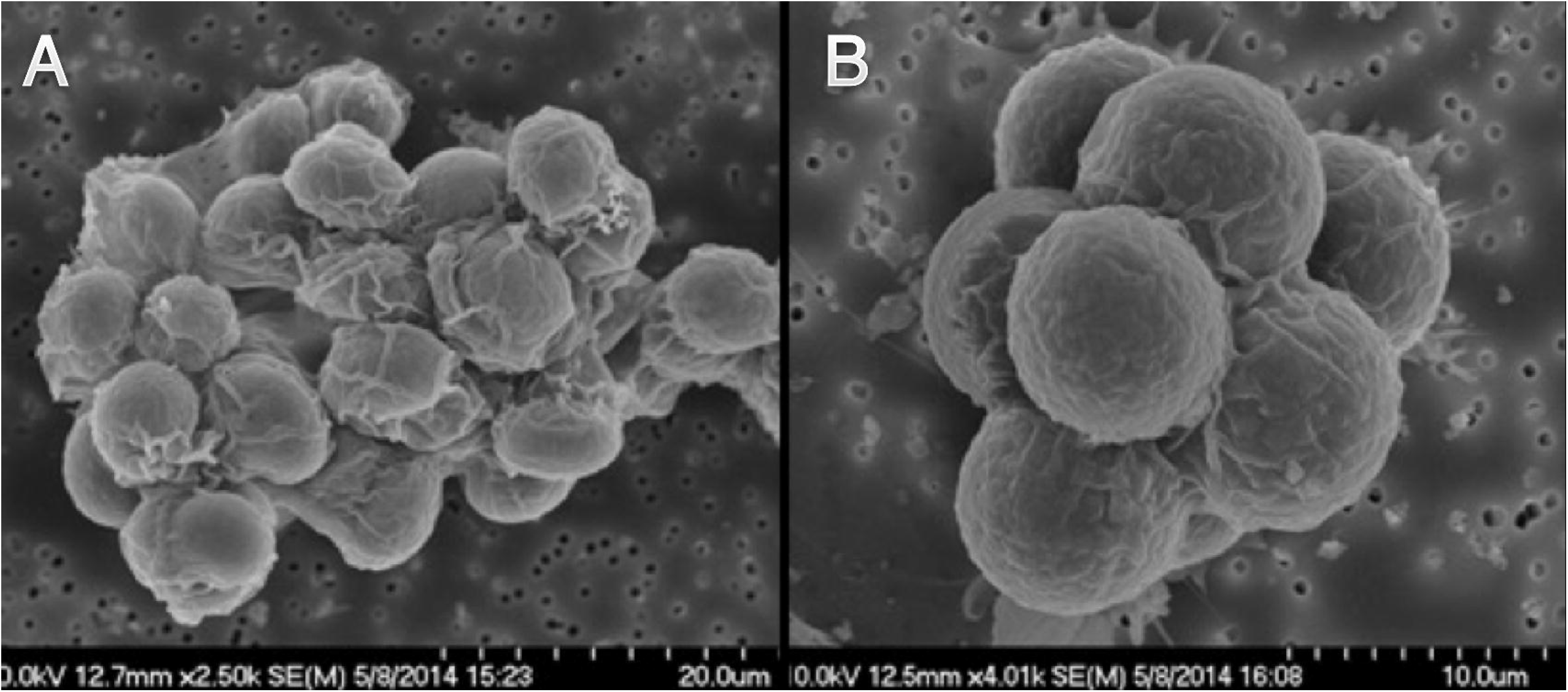

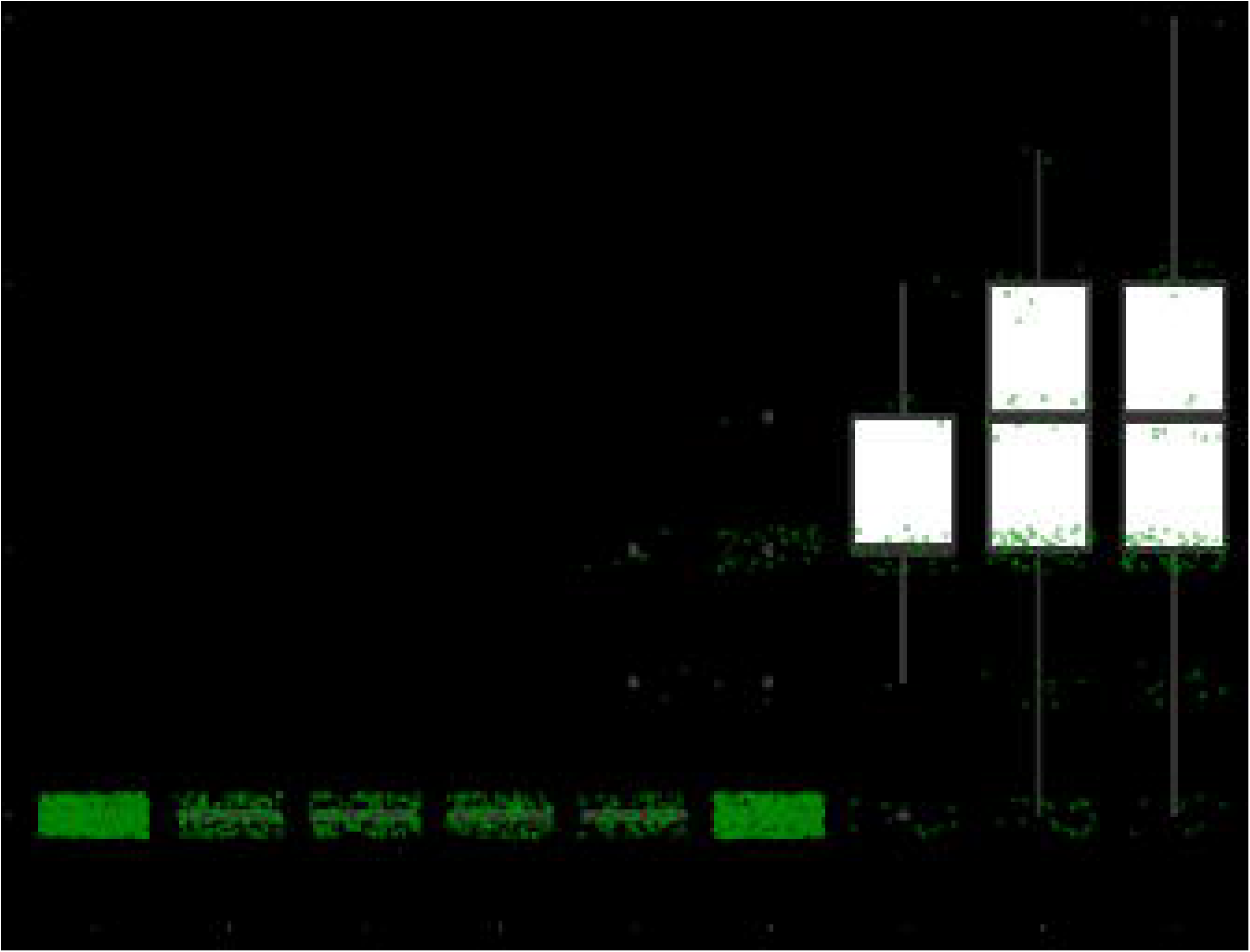

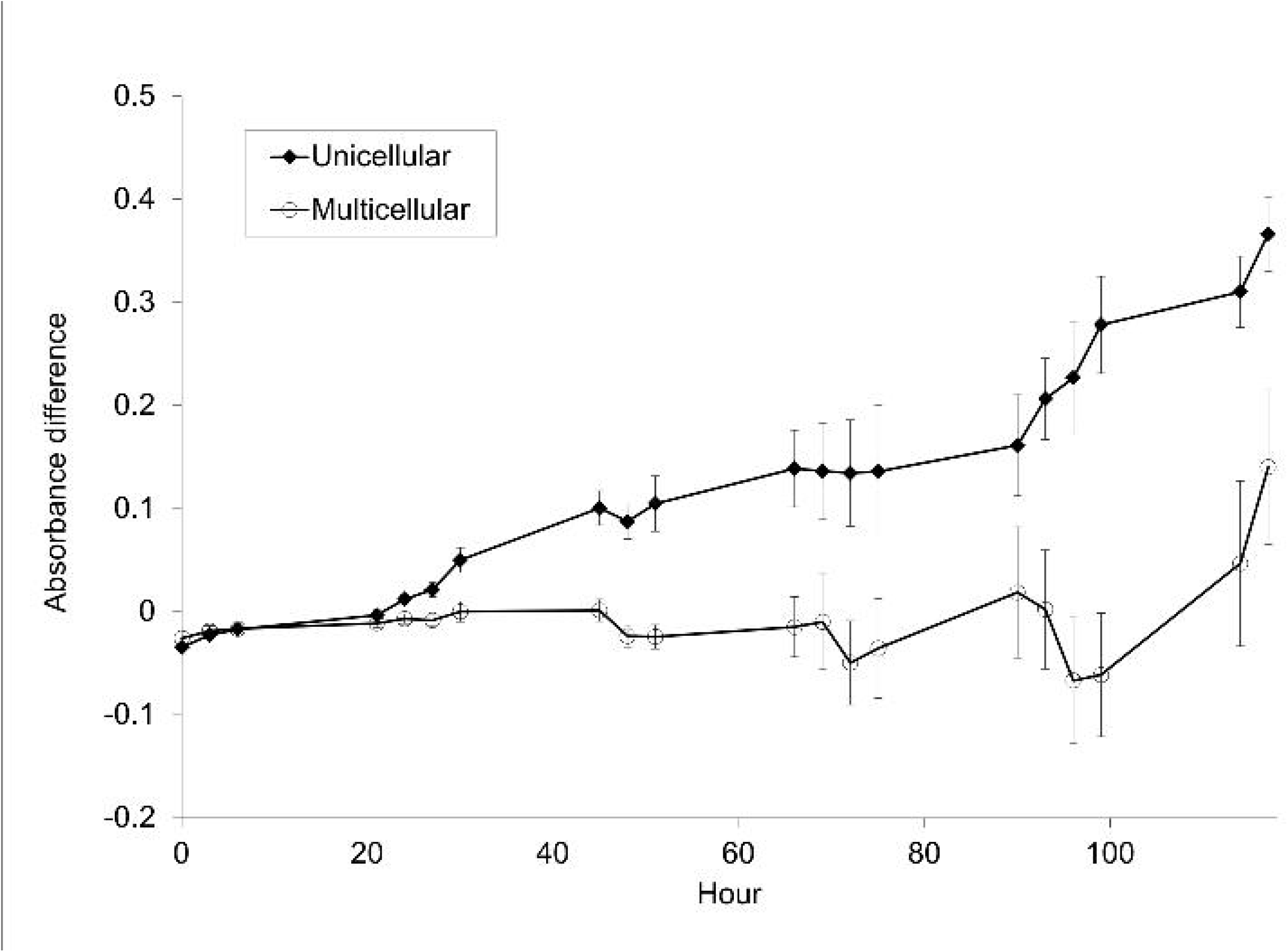

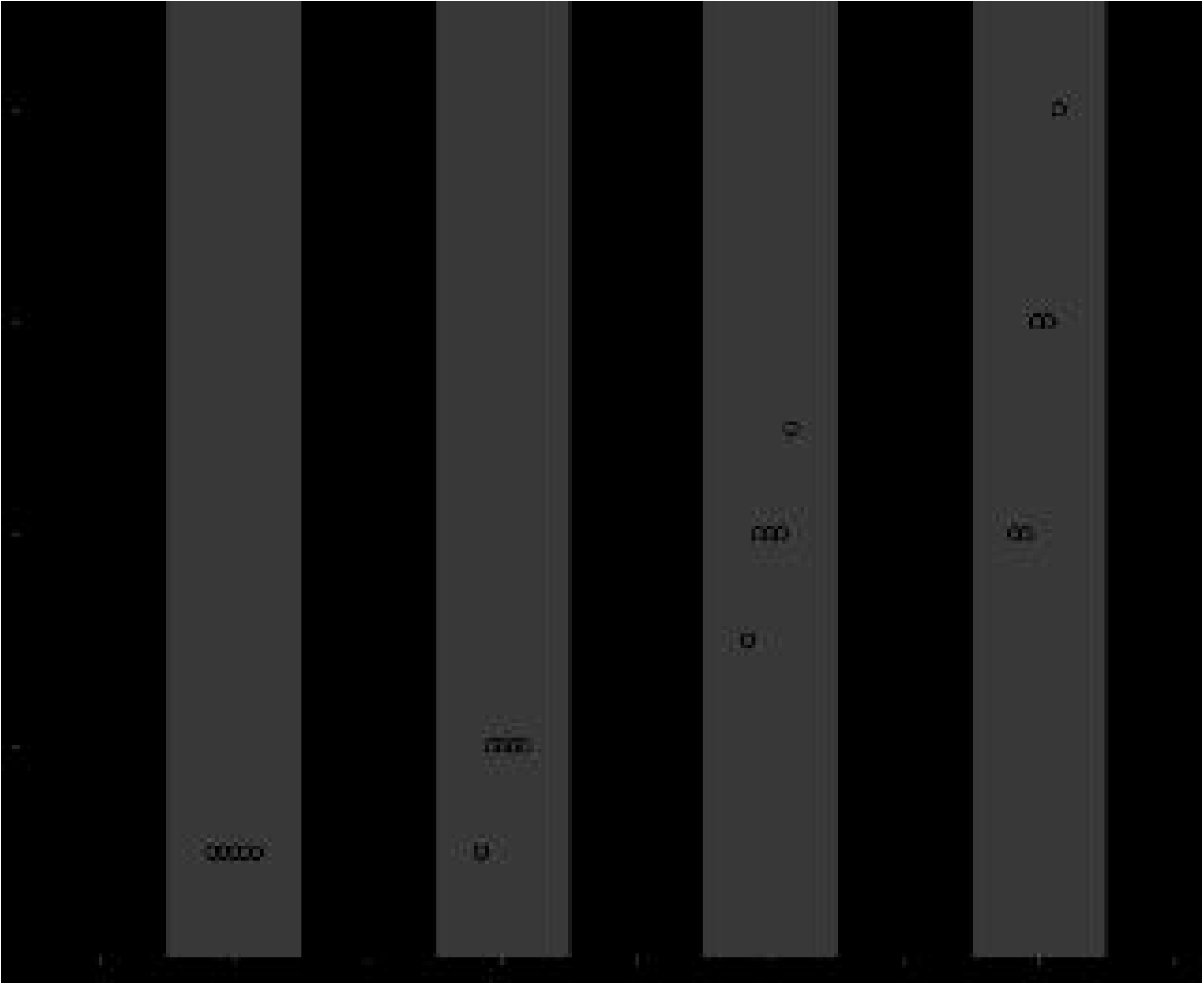

## Acknowledgments

This work was supported by the National Science Foundation (DEB-1457701, DEB-1456652), the John Templeton Foundation (43285), the NASA Astrobiology Institute (NNA17BB05A), a NASA Postdoctoral Program Fellowship to MDH, and a Packard Foundation Fellowship to WCR.

## Methods

### Experimental evolution

To increase genetic diversity at the onset of the experiment, we founded five experimental (B1–B5) and three control (K1–K3) populations from an F1 cross. We obtained plus (CC-1690) and minus (CC-2290) mating type strains of *C. reinhardtii* from the *Chlamydomonas* Resource Center (www.chlamycollection.org) and crossed them using a protocol adapted from Harris^18^. Vegetative cells were grown to high density in 10 mL TAP medium^19^ at 22.5°C and pelleted in 15 mL conical tubes by centrifugation for 5 min. at 2000 × *g*. Pelleted cells were resuspended in 10 mL mating medium (TAP minus NH_4_Cl) and grown in light for 6 h to induce gametogenesis. Gametes were mixed in a 24-well tissue culture plate with 0.5 mL of each mating type, incubated in light for 4 h, and then dried in the dark. Single mating-type controls of 1 mL were treated identically to ensure that no vegetative material survived desiccation. After 20 days in the dark, crosses and single mating-type controls were each flooded with 1 mL TAP medium and placed in light at 22.5°C to germinate. Single mating-type controls showed no evidence of vegetative growth after two weeks; thus, desiccation was effective in killing vegetative material and surviving algae from the crosses must be outcrossed.

Experimental populations were started by mixing 0.1 mL of *C. reinhardtii* F1 cultures, grown to ~2 × 10^6^ cells/mL, with 0.1 mL of a culture of *P. tetraurelia*, grown to ~2 × 10^5^ cells/mL, in 1.3 mL of COMBO medium^20^, making a final volume of 1.5 mL. Control populations were founded by inoculating 0.1 mL of *C. reinhardtii* F1 cultures in 1.4 mL of COMBO medium. All populations were propagated in 24-well plates and transferred weekly.

During transfers, populations were homogenized with a multichannel pipette by drawing and dispensing 1 mL three times. After mixing, 0.1 mL of culture was transferred to 1.4 mL fresh COMBO in a new 24-well plate. After 350 days, populations were plated on TAP medium + 1.5% agar. For each population plate, eight individual colonies were randomly picked and re-plated on agar. Individual colonies were picked and re-plated at least three times each to ensure that each was monoclonal and free of contaminants.

We initially characterized evolved phenotypes using light microscopy. A homogenized dilution of each isolate grown in TAP medium was fixed in iodine and stored at 4°C. For each isolate, 48 μL samples were photographed in replicate on a hemocytometer. Images for each of the clones were processed in ImageJ to determine particle counts and relative particle sizes, and these data were later processed in Python to determine the relative proportions of different sized particles. The size threshold used to distinguish between unicellular and multicellular particles was user-determined and was rarely violated by any samples. In all instances where the size threshold was violated, small multicellular particles were categorized as unicellular; thus, we remain conservative in our characterization of particle sizes, under- rather than overestimating the number of cells in particles.

### Maximum Cells Per Cluster

In order to determine the size attained by different evolved strains, measured as the maximum number of cells per cluster, we stained and imaged samples via fluorescent microscopy over a 6-day examination period. Strains were cultured in 10 mL of TAP medium in a climate-controlled chamber on a 14:10-hour light:dark cycle (23.5°C, 41% humidity, 326 lx). In order to standardize starting cell concentrations, preliminary cultures were first inoculated from scrapes of colonies on TAP plates (2% agar) and allowed to grow, without disruption, for 6 days until reaching stationary phase. Cultures were then transferred 1:100 into fresh TAP medium and examined over the following 6-day examination period.

Throughout the examination period, cultures were sampled, stained, and imaged every 12 hours in order to capture the maximum size attained by various strains before reaching stationary phase. Strains B5-05 and B5-06 were not sampled at the final time-point. Samples were extracted from well-mixed culture tubes and transferred into 1.5 mL microcentrifuge tubes (200 μL from the first 6 time-points to compensate for low cell density, 100 μL from all time-points thereafter). The samples were centrifuged at 250 x *g* for 5 minutes and the supernatant was replaced by a 50% v/v ethanol-water solution to fix cells and make cell membranes more permeable to the DAPI (4’,6-diamidino-2-phenylindole) fluorescent nucleotide stain. After a 10-minute fixation period, samples were centrifuged again (250 x *g* for 5 minutes) and the supernatant was replaced with distilled water containing 1 μg/mL DAPI. Cells with DAPI were kept in darkness for one hour and then moved to a refrigerator without light to prevent photo bleaching until imaging on the microscope.

Prior to imaging, samples were removed from the refrigerator and inverted to mix cultures. Seven μL of each sample were mounted on glass microscope slides and large 6 × 6 images were stitched from individual images captured at 40x magnification, creating high-resolution (441 megapixel) wide-field images, providing a robust measurement of the distribution of clusters sizes within the population.

### Image and data analysis

Large stitched images of individual samples were imported and analyzed in Fiji^21^. We wrote and implemented a script in ImageJ macro (Supplemental Material) to automatically calculate the number of cells in each cluster. The script accomplished this by demarcating cluster boundaries and recording the number of pixel maxima within each boundary, corresponding to the number of DAPI-stained nuclei. Following this procedure, all images used in our analyses were manually screened and any regions of interest resulting from artifacts (*e.g*., dust particles, autofluorescence) were removed from the dataset. Data for five technical replicates of all tested strains, each represented by 12 time-points sampled over six days, were concatenated into one large dataset.

In order to determine the maximum number of cells per cluster achieved by each strain, hereafter referred to as “cluster size”, we first determined the time-point at which clusters were largest. This was calculated separately for each strain, as different strains reached their maximum size at different time-points during the six-day culture cycle. First, we calculated the median cluster size of each replicate at each time-point throughout the culture cycle. Medians were chosen to represent the central tendency instead of the mean because the distributions of cluster sizes were right-skewed and occasionally contained large outlier clusters. Then, to determine the time-point at which cluster size was largest for each strain, we calculated the average (mean) of the median cluster sizes from replicate populations within each time-point. The first time-point, representing cluster sizes at time of inoculation, was removed from analysis.

### Number of Cells per Propagule

In order to determine the number of cells per propagule released by clusters of each strain, hereafter referred to as “propagule size”, we time-lapse imaged all strains over 72 hours of growth. First, cells were inoculated from colonies on TAP agar plates into liquid TAP media and incubated for five days to grow to high density. Then, each culture was homogenized by vortexing, transferred 1:100 into five replicate tubes of fresh TAP media, and allowed to grow for two days. These exponentially growing cultures were mixed by vortexing, diluted 1:10 in TAP medium, and then 100 μL from each culture was randomly inoculated into the central 60 wells of a 96-well tissue culture plate that already contained 100 μL of TAP medium in each well.

For time-lapse microscopy, the 96-well plate (containing five technical replicates of each strain) was imaged at 200x magnification, where the field of view was positioned on cells near the center and at the bottom of wells. The time-lapse was run for 72 hours, capturing images of each well every 30 minutes.

Time-lapse images were manually analyzed using ImageJ. For each well, all images from the time-lapse were viewed individually and sequentially. All cells or clusters of cells in the frame at the beginning of the time-lapse were labeled with a number indicating their identity. Throughout the time-lapse, when a cell or cluster of cells was seen to separate from the initial parental cluster/group, the number of cells in the propagule was recorded. For wells that had few or no initial cells in the frame (typically occurring for motile, unicellular strains), cells or clusters of cells were labeled as they entered the field of view, until collecting a sufficient sample size (N > 5 parental clusters). Due to the long time required for clusters to fully separate physically—even after they have detached from a parent cluster—propagule sizes were recorded four frames (two hours) after the propagule was observed to initially split from the parental cluster. Data collected from time-lapse images were analyzed in R^22^.

### Defense against predators

To test whether multicellularity affords protection from predators, we subjected evolved unicellular and multicellular isolates to predation by the rotifer *Brachionus calyciflorus*. Rotifers were chosen for this assay instead of *Paramecium* to reduce the likelihood that some adaptation other than cluster formation protected cluster-forming strains from the predator. Rotifers were obtained as cysts in vials and stored at 4°C in the dark. To ensure that cultures of *B. calyciflorus* were free of contaminants, cysts were treated and incubated according to the protocol described by Suga *et al*.^23^. Neonate rotifers were grown axenically in 10 mL of WC medium^24^ with 100 μg/mL ampicillin in culture tubes at the same temperature and light conditions as the *C. reinhardtii* cultures. In order to sustain the population of *B. calyciflorus* before the experiment, 1.0 mL of an axenic, unicellular strain of *C. reinhardtii*, CC-1690, was inoculated into the culture via micropipette.

Evolved isolates of *C. reinhardtii* were grown to high density, and culture absorbance was measured using a microplate reader (Molecular Devices SpectraMax^®^ M5, 420 nm), whereafter each was diluted to approximately 3.45 × 10^5^ cells/mL. Four replicates of each strain were randomly assigned to 24-well plates and 1.33 mL of each strain pipetted into designated wells. Twelve multicellular strains (B2-03, 04, 06, 07, and 11; B5-01 through 07) and twelve unicellular strains (B2-01, 10, and 12; B5-08; K1-01, 02, and 04 through 09) were used for this assay. Rotifer cultures were poured through a 25 μm mesh filter to remove the algae they had been feeding on, then pipetted into a sterile test tube awaiting transfer to the experimental wells (*C. reinhardtii* cells are ~10 μm in diameter and easily pass through the filter). Depending on the treatment condition, 0.67 mL of either WC medium or WC medium with predators was added to each well. Initial absorbance values were recorded and then plates were stored under lights in an incubator on a 14/10-hour light/dark cycle at 22.5 °C for 5 days. Throughout the 14-hour light period, absorbance readings were taken every 3 hours, providing growth rates with high resolution for all strains.

